# Rapid and sensitive detection of genome contamination at scale with FCS-GX

**DOI:** 10.1101/2023.06.02.543519

**Authors:** Alexander Astashyn, Eric S. Tvedte, Deacon Sweeney, Victor Sapojnikov, Nathan Bouk, Victor Joukov, Eyal Mozes, Pooja K. Strope, Pape M. Sylla, Lukas Wagner, Shelby L. Bidwell, Karen Clark, Emily W. Davis, Brian Smith-White, Wratko Hlavina, Kim D. Pruitt, Valerie A. Schneider, Terence D. Murphy

**Affiliations:** National Center for Biotechnology Information, National Library of Medicine, National Institutes of Health, Bethesda, MD, USA

**Author notes:** A.A. and E.S.T. contributed equally to this work. All authors: National Center for Biotechnology Information, National Library of Medicine, National Institutes of Health, Bethesda, MD, USA.

**Keywords:** Genome contamination, Genome quality, Genome assembly, GenBank, RefSeq, Software

## Abstract

Assembled genome sequences are being generated at an exponential rate. Here we present FCS-GX, part of NCBI’s Foreign Contamination Screen (FCS) tool suite, optimized to identify and remove contaminant sequences in new genomes. FCS-GX screens most genomes in 0.1-10 minutes. Testing FCS-GX on artificially fragmented genomes demonstrates sensitivity >95% for diverse contaminant species and specificity >99.93%. We used FCS-GX to screen 1.6 million GenBank assemblies and identified 36.8 Gbp of contamination (0.16% of total bases), with half from 161 assemblies. We updated assemblies in NCBI RefSeq to reduce detected contamination to 0.01% of bases. FCS-GX is available at https://github.com/ncbi/fcs/.

## Background

The National Center for Biotechnology Information (NCBI) currently hosts over one-and-a-half million genome assemblies submitted to the archival GenBank, European Nucleotide Archive (ENA), and DNA Data Bank of Japan (DDBJ) databases of the International Nucleotide Sequence Database Collaboration (INSDC), commonly referred to as “GenBank”, totaling >22 terabases of genome sequence data. Decreased sequencing costs have accelerated the production of genome assemblies and their submission to public databases, with sequence bases in GenBank doubling around every 18 months [1]. While all sequences submitted as part of a genome assembly should originate from the declared source organism, a subset of sequences often contains foreign DNA as genome contamination. Contamination can occur at multiple stages of a genome assembly project [2] and can arise from challenges intrinsic to an organism’s biology, such as other organisms in the surrounding environment or the presence of endosymbionts [3]. New sources of contamination have been introduced following advances in genomics methods including multiplexing [4, 5] and metagenome-assembled genomes (MAGs) [6]. High-quality genomes are essential for data analysis across biological disciplines, whereas contamination confounds biological inference. Contaminated sequences have formed the basis for incorrect conclusions regarding evolutionary relationships [7] and lateral gene transfer [8]. The problems of contaminants are compounded when misidentified sequences are submitted to public archives. There are numerous reports of contaminants in NCBI databases [9-15], including in the genomes of model organisms [9, 16]. In assuming databases and their associated tools are error-free, researchers performing comparative genomic analysis might be confused by illogical results or publish findings based on artifactual connections. Of particular concern is that the addition of contaminated sequences and associated annotations into databases can perpetuate errors when the databases themselves are used for future annotation efforts, contributing to a vicious cycle [11].

Several genome contamination detection tools have been developed to address this emergent issue [2]. Current tools differ in scope and implementation, such as their assessment of contamination at an assembly level or sequence level, their application for certain taxonomic groups, and their use of reference-based or database-free methods. Moreover, most tools aren’t designed for automated analysis and removal of contaminant sequences from new assemblies. The legacy NCBI contamination detection pipeline screens new genome submissions using VecScreen [17] and BLAST [18]. In 2022, NCBI identified contaminants in one out of every ten prokaryote genomes and one out of every three eukaryote genomes (**Table 1**). However, NCBI pipelines miss a nontrivial amount of contamination, evidenced by the published reports discussed above. Publicly-available contamination screening tools are foundational to support the new NIH Comparative Genomics Resource (CGR) project goal of enabling reliable comparative genomics analyses in eukaryotic research organisms [19].

**Table 1.** Summary of contamination detected in genome submissions to NCBI in 2022.

| Organism group | Total submissions | Contaminated submissions<sup>1</sup> | Common contaminants<sup>2</sup> | Contamination matching other foreign sequences |
| --- | --- | --- | --- | --- |
| Bacteria & Archaea | 155,618 | 16,263 | 9,381 | 8,647 |
| Eukaryota | 11,232 | 4,063 | 3,196 | 2,613 |
| Other genomes | 3,771 | 413 | 338 | 227 |
| Total | 170,621 | 20,739 | 12,925 | 11,487 |

<sup>1</sup>submissions contaminated with common contaminants, other foreign sequences, or both
<sup>2</sup>includes adaptors, vectors, bacterial insertion sequences, and bacteria/virus genomes commonly found in assemblies (e.g. *E. coli*, phiX174)

Here we present GX, a new genome cross-species aligner to identify genome contamination from foreign organisms using hashed k-mer (h-mer) matches and a curated reference database (**Fig. 1**). GX is part of the NCBI Foreign Contamination Screen (FCS) tool suite available at https://github.com/ncbi/fcs. We demonstrate FCS-GX taxonomic identification is highly accurate in simulated genomes from diverse taxa. We additionally characterize contamination in GenBank genomes and reduce detectable contamination in NCBI RefSeq [20]. The source code for FCS-GX is available at https://github.com/ncbi/fcs-gx.

**Fig. 1.**
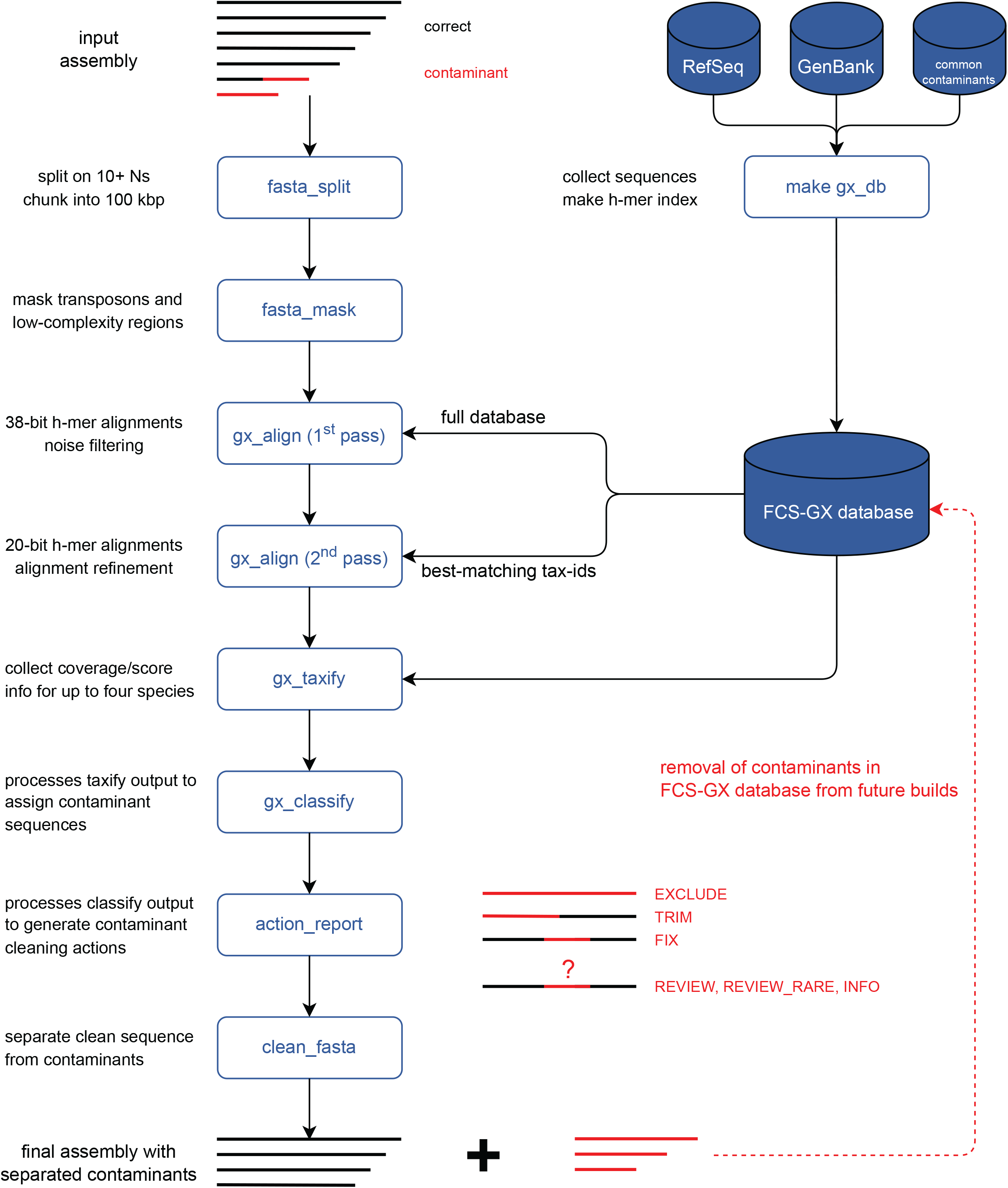
Overview of FCS-GX pipeline. FCS-GX splits genome assembly scaffolds into contigs and chunks contigs into 100 kbp subsequences for processing. FCS-GX performs repeat detection and masking in eukaryote assemblies. The GX aligner operates in two passes using modified k-mers (h-mers) to align query sequences first to the entire indexed reference database and second to sequences corresponding to the tax-id sets providing best matches for alignment refinement. After collecting coverage and score information FCS-GX assigns likely contaminant sequences by comparing the taxonomic assignment calculated for each sequence by the user-specified tax-id. The final output from FCS-GX is a cleaned FASTA alongside an action report that details contaminant cleaning actions taken (FCS-GX actions EXCLUDE, TRIM, FIX) as well as details for additional sequences warranting manual review but are not automatically cleaned (FCS-GX actions REVIEW, REVIEW_RARE, INFO). See Methods for descriptions of FCS-GX action categories. FCS-GX uses a custom reference database totaling 709 Gbp of sequence data from assemblies and common contaminants used in current NCBI screening. Assemblies contributing to the database were screened by FCS-GX while excluding self-hits. High-confidence contaminants were removed in order to use the database for screening new genomes.

## Results

We began development on FCS-GX with the goal of improving sensitivity to contaminants without compromising specificity. *Ad hoc* analyses of known contaminated genomes indicated a need for a large and diverse screening database to detect the diversity of potential contaminants and distinguish them from correct sequences. Furthermore, contaminants may represent novel strains or species, necessitating an approach that does not require high identity alignments.

We designed FCS-GX to address these challenges by identifying potential sequence matches through hashed k-mers (h-mers) which are modified to allow for matches of non-identical sequences. H-mer matches are extended into longer gapped alignments to improve coverage, and intra-genome repeats and low-complexity sequences are identified to reduce false positives. FCS-GX can assign taxonomic labels to sequences by reporting and interpreting alignment score information from one or multiple taxonomic divisions. FCS-GX screens against a diverse reference database of 709 Gbp including assemblies from 47,754 taxa (database build-date 2023-01-24), which is size optimized to fit in the memory of a 512 GiB server.

We prioritized speed and ease-of-use to distribute FCS-GX as a publicly available tool that assembly providers could run early in genome assembly pipelines, resulting in better assemblies and easier submission to NCBI GenBank. Overall execution time is the sum of reading the database into memory, which can take 4-30+ minutes depending on the source and hardware, followed by screening which takes 0.1-10 minutes/genome for most species. FCS-GX requires a user-provided genome in FASTA format along with a NCBI taxonomic identifier (tax-id) [21] as inputs. Identification and removal of contamination in both eukaryote and prokaryote genomes is automated with minimal user interaction (**Additional file 1: Fig. S1**). As such, screening with FCS-GX can accommodate the current exponential growth in genome sequencing.

### FCS-GX accurately detects contaminants with few false positives

High sensitivity and specificity are critical for automated screening and trust in results; false assignment of longer or particular types of sequences as contaminants could lead to loss of substantial content in screened genomes. To measure sensitivity and specificity, we reasoned that long, gap-free sequence spans from highly contiguous genomes are likely to be contaminant free and could be used to test FCS-GX. Since contaminant sequences tend to be short, we artificially fragmented sequences into subsequences of a defined size (1, 10, or 100 kbp), and tested them in two ways. First, we tested sensitivity by running FCS-GX with a discordant species such as human for an alphaproteobacteria genome; a genome where every sequence is identified as alphaproteobacteria reflects a sensitivity value of Sn=100%. Second, we tested specificity by running FCS-GX with the expected species; an alphaproteobacteria genome with no contaminant calls reflects a specificity value of Sp=100%.

FCS-GX exhibited high sensitivity across diverse samples from five tested “kingdom” groups (Metazoa, Viridiplantae, Fungi, other Eukaryotes, prokaryotes) when the contaminating species is in the FCS-GX database (e.g., strain-level differences): 76% of prokaryote and 91% of eukaryote datasets achieved better than Sn=95% with 1 kbp fragments across 663 and 370 species assayed, respectively, with near 100% sensitivity achieved for most species at larger fragment sizes (**Fig. 2A, Additional file 2: Table S1**). Reduced sensitivity in small sequences is generally due to poor alignment coverage or inconclusive taxonomic assignments resulting in non-contaminant calls and not due to incorrect taxonomy assignments (**Additional file 1: Fig. S2, Additional file 2: Table S2**).

**Fig. 2.**
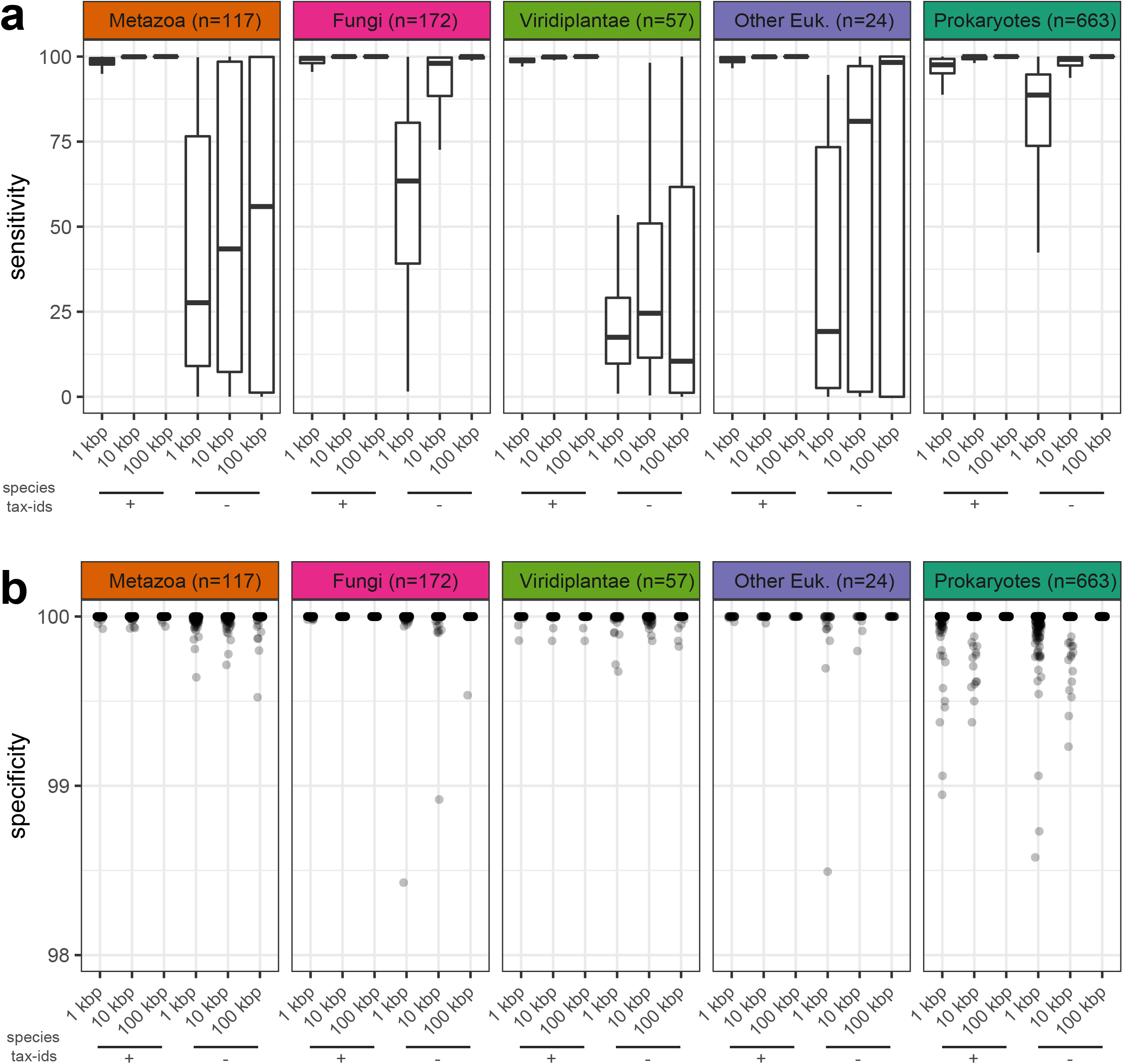
Sensitivity and specificity of FCS-GX contamination detection. **a** Distributions of sensitivity measurements. Distributions are shown for artificially fragmented genomes in five “kingdom” groups. Sensitivity is shown for genomes fragmented at three different window sizes (1 kbp, 10 kbp, 100 kbp). For each window size, sensitivity is shown for FCS-GX runs while including the same species tax-ids as the source genome during the alignment stage (+) and while excluding same species tax-ids (-). **b** Distributions of specificity measurements for the same set of fragmented genomes in **a**. The dotplot shows an enlarged view of the upper limit of specificity (98-100%). The full dotplot including ten outliers not visualized here is available at **Additional file 1: Fig. S4**. See **Additional file 2: Table S1** for complete sensitivity and specificity score data.

We simulated contamination detection of novel organisms by selectively dropping out sets of tax-ids corresponding to the species of the source organism from the FCS-GX alignment stage as if they did not exist in the reference database. Sensitivity decreased when simulating novel contaminant species, dropping to a median sensitivity of 89% for prokaryote and 17-63% for eukaryote 1 kbp fragments (**Fig. 2A**). We observed a positive association between aggregate genome coverage by FCS-GX alignments and sensitivity (**Additional file 1: Fig. S3**). The larger representation of prokaryotes and Fungi in the FCS-GX database relative to Metazoa, Viridiplantae, and other Eukaryotes contributes to a higher frequency of robust alignment coverage when simulating novel species and results in better Sn scores.

FCS-GX specificity tests indicated a low incidence of false positives, with lower Sp scores observed for smaller fragment sizes alone or in combination with exclusion of species tax-ids, although differences were marginal and not easily visualized (**Fig. 2B, Additional file 1: Fig. S4, Additional file 2: Table S1)**. When manually inspecting lower Sp score outliers we found a mix of valid contaminants assembled in large sequences as well as false positives (**Additional file 2: Table S3**). 95% of prokaryote datasets achieved Sp=100% with 1 kbp fragments, with a marginal decrease to 88% when excluding same-species tax-ids. Most false positives correspond to sequences assigned to other prokaryote taxonomic divisions and are below 1% of total genome length which are not flagged for cleanup in GenBank submission processing (see Methods).

We estimated sequence-level specificity by treating all intra-kingdom contaminants as false positives and subtracting all inter-kingdom contaminants from the remaining sequences to count true negatives. Sp scores were >99.93% in all scenarios, and >99.97% when the same species is in the database (**Additional file 2: Table S4**). Treating all inter- and intra-kingdom contaminants as false positives had little effect on Sp scores (**Additional file 2: Table S4**).

### FCS-GX enables high-throughput contamination screening

After loading the database into memory on a single 64 vCPU server, we completed screens of 28,774 eukaryote genomes totaling 15.7 Tbp in 18 days. We completed screens on batch runs of prokaryote genomes on servers of similar capacity with a net throughput of 1.94 sec/genome. Compared to the legacy screen used for NCBI genome submissions, we calculated that megaBLAST uses 135x more CPU time relative to FCS-GX while aligning against 80% less sequence. Thus, we demonstrate FCS-GX is scalable to high-throughput assembly projects.

### FCS-GX detects extensive contamination in NCBI databases

We characterized contamination in 1,545,312 prokaryote and 30,053 eukaryote genome assemblies totaling 22.4 Tbp of sequence data in the current GenBank archives (April 15, 2023). We identified 36.8 Gbp of suspected contamination from 23,451,091 sequences, equivalent to 0.16% of the total bases and 1.30% of the sequences assayed, and including 2,932,319 annotated proteins (**Additional file 2: Table S5**). The distribution of the proportion of contaminated sequence per genome was bimodal with peaks approaching the 0% and 100% extremes (**Fig. 3A**). The total length of contaminated sequence has increased along with the total length of GenBank genomes over time (**Fig. 3B**) such that the percentage of contaminated sequence has remained steady over time (**Fig. 3C, Additional file 2: Table S6**).

**Fig. 3.**
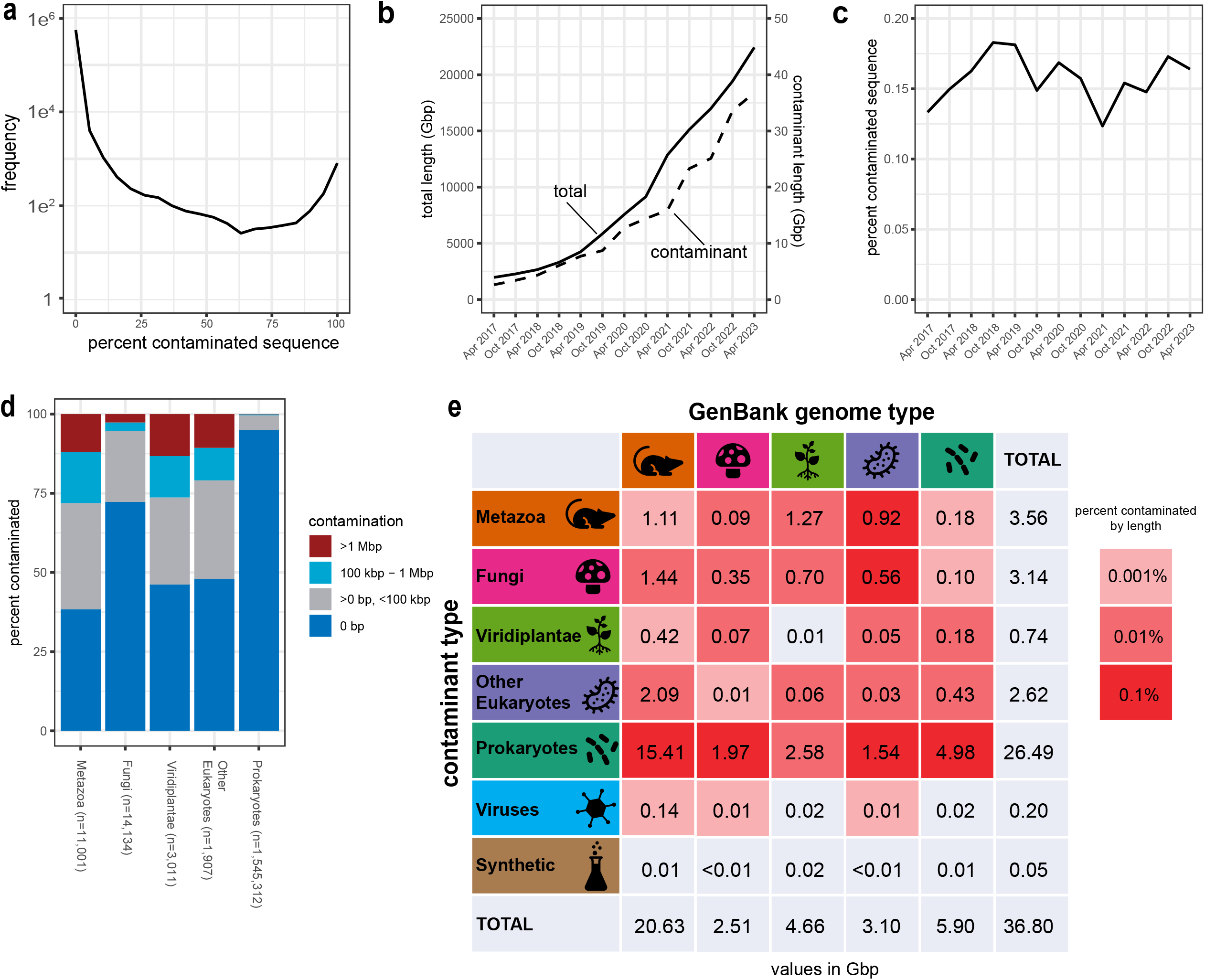
FCS-GX detection of contamination in NCBI databases. **a** Distribution of the proportion of contaminated sequence per genome detected by FCS-GX in the NCBI GenBank database. Genome counts (frequency) were computed in 5% intervals. **b** Aggregate length of total genome sequence (solid line) and contaminated sequence detected by FCS-GX (dashed line) in the NCBI GenBank database from 2017 to 2023. **c** Percentage of contaminated sequence detected by FCS-GX (dashed line) in the NCBI GenBank database from 2017 to 2023, *i.e.,* the quotient of the contaminant amount divided by the total amount displayed in **b**. See **Additional file 2: Table S6** for supporting numerical data. **D** Percentage of contaminated genomes in GenBank and RefSeq. Total numbers of screened genomes are shown for five taxonomic “kingdom” groups: Metazoa (animals), Fungi, Viridiplantae (green plants), Other eukaryotes, and Prokaryotes (Bacteria + Archaea). Within each group, genomes are placed into four bins corresponding to the amount of contamination per genome and percentages are calculated for the count of genomes in each bin divided by total screened genomes. **E** Aggregate contamination lengths identified in genomes from five “kingdom” groups. Colors of grid squares indicate aggregate contamination lengths from seven sources (five kingdoms, plus virus and synthetic) that correspond to percentages of total assembly length for each GenBank kingdom group. See **Additional file 2: Table S5** for supporting numerical kingdom contamination summary data.

However, contaminants are not evenly distributed: the representative genome set, which includes only one genome per species, was 74% cleaner than non-representative genomes (0.056% vs 0.215% by length), prokaryote genomes from multiisolate projects focused on sequencing multiple genomes for a single species were 98% cleaner than other prokaryote genomes (0.007% vs 0.312%), and genomes with high contiguity that are often the product of long read sequencing (contig N50 >1 Mbp) were 87% cleaner than other genomes (0.028% vs 0.209%). Contaminant sequences were typically small: 82% of contaminants were ≤1 kbp and 98% of contaminants were ≤10 kbp (**Additional file 1: Fig. S5**). Since short sequences are rarely annotated, we added an option to drop all sequences below a size threshold and recommend using a 1 kbp threshold for eukaryote genomes.

Next we assessed contamination patterns across multiple taxonomic ranks. The contamination rate was lower in prokaryotes relative to eukaryotes with Fungi genomes having lower contamination rates and a lower percentage of heavily contaminated genomes (>1 Mbp) relative to Metazoa, Viridiplantae, and other Eukaryotes (**Fig. 3D**). Prokaryote contaminants represent 26.2 Gbp (71%) of the aggregate contamination, including 15.4 Gbp found in Metazoan genomes (**Fig. 3E, Additional file 2: Table S5**). When looking at genomes grouped into taxonomic divisions used by FCS-GX to assign sequences, we found examples of pervasive contamination with obvious biological connections, such as alveolate-in-bird, alphaproteobacteria-in-insect, and insect-in-plant (**Additional file 2: Table S7**).

### Biological sources of contamination

Genome contamination often reflects the underlying biology of the organism and can derive from symbionts, infection, gut and surface microbes, and diet. Symbionts and parasites are common contaminants when sequencing host genomes or environmental samples. We found 864.4 Mbp of contamination with the apicomplexan parasite *Sarcocystis neurona* as the top hit from the FCS-GX database, mostly in mammal and bird genomes. Authors of the contamination detection software BlobToolKit previously identified a bird genome (*Crypturellus cinnamomeus*; GCA_003342915.1) with apparent contamination from a whale (*Physeter catodon*; GCF_002837175.2) that they ultimately explained as shared *Sarcocystis*-like contamination in both genomes [22]. FCS-GX directly confirmed *Sarcocystis* contamination in both, including 130 Mbp in *C. cinnamomeus*, and found an additional 24.3 Mbp in *P. catodon* with a 13.8 Mbp contaminant span at the beginning of the 145.7 Mbp chromosome 1 sequence. The reverse can also be true where host DNA sequences are assembled alongside the genomes of parasites, as was the case of 2.0 Mbp fish sequence identified in the salmon louse *Lepeophtheirus salmonis* (GCA_016086655.1). We identified examples of symbiont contaminants across diverse samples where the FCS-GX database had closely related species to the suspected contaminant: green algae in a fungal component of a lichen (*Cladonia squamosa*; GCA_947623385.1), fungi in an insect (*Nilaparvata lugens*; GCA_014356525.1), and protist in a prokaryote MAG produced from an environment sample (*Cohaesibacter* sp.; GCA_025800105.1). FCS-GX can also detect novel biological contaminants, such as 2.2 Mbp of an unknown CFB group bacteria in two haplotype assemblies of a toad (*Spea bombifrons;* GCA_027358695.1, GCA_027382365.1) and 359.4 kbp of an unknown bacteria in a fly (*Condylostylus longicornis*; GCA_029603195.1).

### Experimental sources of contamination

The recent growth of community projects aimed at producing reference genomes for taxonomic groups and geographical areas has contributed to quality improvements of genome data in public databases. Even so, the sequencing and assembly of diverse samples can be a source of cross-contamination. We found 812.9 Mbp contamination most similar to the dinoflagellate coral symbiont *Cladocopium goreaui*. 803.3 Mbp of identified *C. goreaui*-like contaminants were in the genomes of basal metazoans submitted by a single submitter (**Additional file 2: Table S8**). We also found examples of genome assemblies from birds, insects, and plants from the same submitter that had >100 kbp *C. goreaui*-like sequences. Since these diverse genomes were made publicly available in the same timeframe as the basal metazoan assemblies (2022-2023), we interpret these patterns as cross-sample contamination that likely arose during sample preparation and/or genome sequencing. We found additional examples of experimental contamination: 200 Mbp from an eagle (*Haliaeetus sp.*) in other animal and plant genomes, 10 Mbp from maize (*Zea mays*) traced back to the use of cornstarch as an absorbant in sample shipping, and 5 Mbp from a moth (*Bombyx mori*) in gammaproteobacterial genomes.

### Extreme genome contamination

We found half of overall contamination in current databases originates from only 161 genomes (range: 32 to 4,512 Mbp, **Additional file 2: Table S9**). These genomes are highly fragmented with suspected contaminants occurring predominantly in small sequences: 95 genomes have a contig N50 <10 kbp and 146 have a contig N50 <100kbp. In addition, we found 1,040 genomes have extreme contamination by proportion of contaminated sequence (90-100%, **Fig. 3A**, **Additional file 2: Table S10**). We expected that some cases would be caused by issues with species assignments associated with the genomes.

Consistent with this prediction, we found genomes that had contaminants reported from a lower taxonomic division, e.g. gammaproteobacterial contamination in a genome declared in the metadata as a more generic proteobacteria, indicating FCS-GX can be used to help improve taxonomic assignment. After examining genomes assigned as heavily contaminated sequence from a more specific taxonomic division, we estimate that half of reported prokaryote-in-prokaryote (2.3 Gbp), Metazoa-in-Other Eukaryote (604.0 Mbp), and Fungi-in-Other Eukaryote (247.7 Mbp) contamination are in scope for taxonomic improvement (**Additional file 2: Table S11**). We identified 37 prokaryote genomes totaling 73.6 Mbp with the reverse situation where generic taxonomic entries in the FCS-GX database caused false positive contamination calls (**Additional file 2: Table S11**). The remainder of extreme contamination cases are typically genomes from various sources identified as nearly entirely prokaryotic sequence (**Additional file 2: Table S10**). In rare cases the submitter comments associated with assembly sequences supports the contamination assignment and may help to correct errors. For example, several insect genomes with extreme prokaryote contamination (e.g. GCA_913698155.1, GCA_913698365.1, GCA_913698315.1) are described in the comments as metagenomically assembled *Rickettsia* genomes. When considering all remaining cases with ≥90% contaminated sequence as misclassified, 233 genomes need taxonomic improvement accounting for 1.0 Gbp of contamination.

### Comparison of contamination detected by FCS-GX vs other methods

Assemblies submitted to NCBI GenBank have historically been screened with a megaBLAST-based process relying on high identity alignments to chromosomes from distant taxa to identify and remove contaminants. We leveraged results from the legacy screen during development of FCS-GX, identifying >98% of known contaminant sequences in a test set of heavily contaminated genomes in addition to novel contaminants due to the increase in sensitivity. To estimate the sensitivity increase, we compared FCS-GX results to the original submission screening data for 14,344 eukaryote and 194,995 prokaryote genomes released in the last 2.5 years, excluding 198 with incorrect or sub-optimal taxonomic information. The contamination by length detected by FCS-GX was 0.163%, representing a fourfold increase in sensitivity over the 0.038% detected by the legacy screen. FCS-GX sensitivity increases are expected given its larger screening database, cross-species alignment method, and ability to detect intra-kingdom contaminants.

Most existing screening methods produce results that either require further manual review (e.g., BlobToolKit [22]) or have taxonomic ranges that aren’t readily compared to FCS-GX (e.g., Physeter [10]). Conterminator uses an all-by-all screening approach that identified over 2 million contaminant sequences in GenBank in 2019 [9]. We identified 16,232 GenBank and 7,023 RefSeq assemblies with one or more contaminant sequences according to Conterminator [9] and compared to FCS-GX. For both sets, FCS-GX confirmed 88% of the suspect sequences identified by Conterminator and found an additional 138-146% of sequences (**Additional file 2: Table S12**). *Ad hoc* review found sequences identified only by Conterminator represent a mix of true and false positives; for example, a 2009 mouse assembly (GCA_000002165.1) had 22,013 sequences flagged as contaminants by Conterminator based on alignment to sequences that are themselves contaminants such as KL772705.1, a plant sequence contaminated by mouse satellite. Half (49%) of the additional sequences found in RefSeq were from intra-kingdom contaminants not in scope for detection by Conterminator, whereas most (80%) of the additional GenBank sequences were cross-kingdom contaminants and represent higher sensitivity by FCS-GX. This is likely explained by the use of conventional k-mers for initial matching and a 90% sequence identity threshold in Conterminator, whereas FCS-GX uses h-mers and relies on score thresholds for filtering which enables detection of novel contaminants through cross-species alignments. Compared to the 2019 Conterminator results, FCS-GX expands the amount of identified contamination in GenBank by sixfold and can be readily applied to screen future individual genomes as they are generated or submitted.

### Cleanup of RefSeq genomes

The INSDC databases provide an archival record and sequences can only be changed or removed with the permission of the submitters. To provide NCBI users with a cleaner subset of genomes, we prioritized the NCBI-curated RefSeq genome collection [20] for contamination cleanup using FCS-GX. We manually reviewed FCS-GX results and used a combination of approaches for an initial round of cleanup on the most heavily contaminated genomes: 1) replacement with newer, higher quality assemblies, 2) removal of contaminated assemblies, or 3) outreach to genome submitters followed by suppression of contaminant sequences in both GenBank and RefSeq and release of an updated assembly version. We also added FCS-GX as a screen before adding new eukaryote genomes into the RefSeq collection, and are in the process of doing the same for prokaryotes. We cleaned 124 eukaryote genomes (**Additional file 2: Table S13**), removing 79,593 sequences totaling 548 Mbp of contamination, including 34,337 genes and 30,356 proteins annotated on contaminant sequences. We previously identified and removed 5,694 suspect prokaryote genomes from the RefSeq collection using ANI [23]; we identified and removed an additional 1,284 genomes using FCS-GX. The current RefSeq collection contains 283,221 prokaryote and 1,616 eukaryote genomes and has 265.1 Mbp of suspected contamination remaining after initial cleanup (**Additional file 2: Table S14**). Contaminated sequence is equivalent to 0.018% of the total prokaryote sequence and 0.003% of the total eukaryote sequence, providing additional support of high FCS-GX specificity (**Fig. 4A**, **Fig 4B, Additional file 2: Table S6**). Overall, we’ve reduced contaminant bases in RefSeq eukaryote and prokaryote genomes by 90% and 53%, respectively, compared to their peaks in 2020 (**Fig. 4A**), and 98% and 81% lower, respectively, compared to GenBank genomes as a whole.

**Fig. 4.**
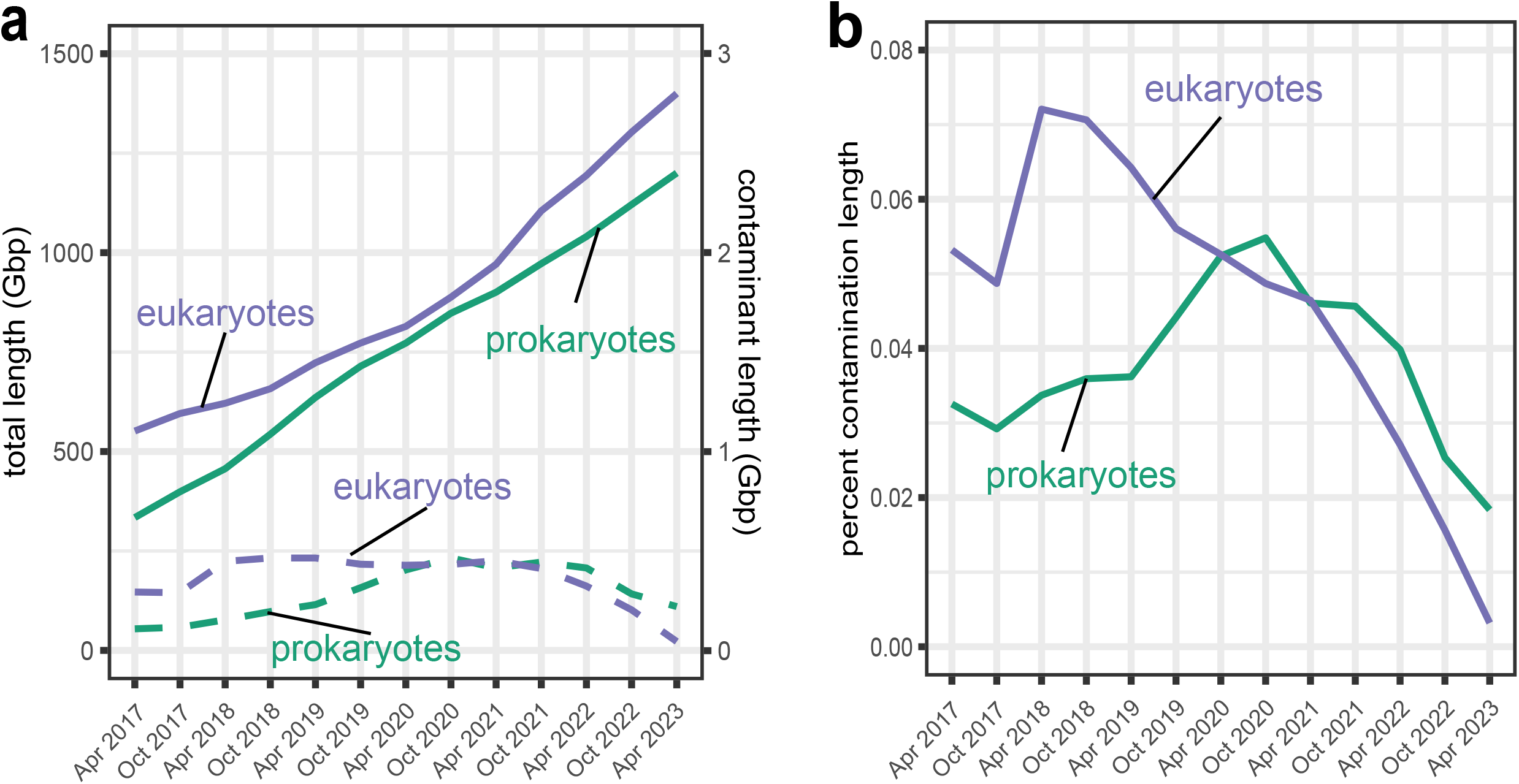
FCS-GX detection of contamination in the NCBI RefSeq database. **A** Aggregate length of total genome sequence (solid line) and contaminated sequence detected by FCS-GX (dashed line) in NCBI RefSeq from 2017 to 2023. **B** Contaminant fraction detected by FCS-GX (dashed line) in NCBI RefSeq database from 2017 to 2023, *i.e.,* the quotient of the contaminant amount divided by the total amount displayed in **a**. See **Additional file 2: Table S14** for supporting numerical Refseq contamination summary data.

### FCS-GX contamination detection is not adversely affected by lateral gene transfer

The transmission of genetic material between genomes via lateral gene transfer (LGT) could result in the improper assignment of transferred sequences as contaminants. Since FCS-GX can classify chimeric sequences with a mix of correct and contaminant spans, we assessed whether FCS-GX reports an excess of chimeric cleaning actions in genomes harboring high-confidence LGTs. We collected FCS-GX results for 3,986 genomes included in the proGenomes2 database with LGT events at the taxonomic family level or above [24]. FCS-GX identified prokaryote-in-prokaryote chimeras in 0.002% (8/3,920) of tested genomes, a similarly low rate compared to 0.004% (2,227/534,869) of chimeras in the current GenBank set excluding multiisolates. We manually inspected the chimeric calls in six genomes with overlaps between the FCS-GX contaminant range and the candidate LGT region. BLAST searches supported the LGT events, but all FCS-GX calls in these regions would not be automatically cleaned (see Methods: FCS-GX output, **Additional file 2: Table S15**). Therefore, we conclude that LGT does not have a systematic confounding effect on FCS-GX contamination detection performance.

Since prokaryote-to-prokaryote LGT is common, the current public version of FCS-GX requires at least a 10 kb span to report a potential chimeric sequence involving contamination from the same kingdom and categorizes them to prompt further analysis before removal. Although prokaryote-to-eukaryote LGT is less common, there are known examples of integrants from endosymbionts affecting multiple host genomes in certain taxonomic groups, such as *Wolbachia* integrants in insects and nematodes [25, 26]. The current public version of FCS-GX implements a special assignment to chimeric sequences involving known sources of LGT (see Methods: FCS-GX output). We found 1176 potential integrants across 366 invertebrate genomes in the current analysis.

## Discussion

We present FCS-GX, a new contamination detection tool to detect foreign sequences in genome assemblies from known and novel organisms. With its fast runtime, high accuracy, and automated removal of contaminants as core features, we recommend that genome submitters use FCS-GX to screen sequences early in the assembly process, such as after the contig assembly stage.

We screened nearly 2 million genomes to assess current and historical contamination in NCBI databases. We identified suspected contamination in genomes representing 0.16% and 0.01% of the genome sequence currently in GenBank and RefSeq, respectively. Most detected contaminant sequences are small and correspond to prokaryote sources. Short eukaryotic genome sequences rarely contribute useful annotation, so we recommend removing sequences below 1 kbp to further reduce contamination levels. Importantly, the proportion of contaminated sequence has not improved over time despite recent advances in sequencing technologies and the availability of new assembly and contamination detection algorithms, reinforcing the urgency for contamination analysis as a standard method in assembly evaluation. The small amount of contamination in current RefSeq reflects the higher stringency for inclusion in the database as well as the cleaning efforts described here. Although we prioritized RefSeq cleanup, we also cleaned 100 GenBank genomes and flagged 409 GenBank genomes with more than 10 Mbp of contamination as ‘contaminated’ in NCBI’s assembly resources to alert users of the data issues.

We demonstrate FCS-GX has higher sensitivity to diverse contaminants compared to other methods. The reduced sensitivity observed for eukaryote divisions not well represented in the FCS-GX database reflects compromises made in database content to keep the size accessible for institutional compute clusters or inexpensive virtual machines from commercial cloud providers, combined with the sparse distribution of coding sequences in most eukaryotes which provide the most signal in cross-species alignments. The database contains genomes from 20,301 prokaryote, 2,419 fungi, and 454 protist taxa, so we expect most environmental contaminants to be detected with very high sensitivity. However, laboratory sources of contamination can include any species, which may ultimately require a substantially larger database for maximal sensitivity. There is also a trade-off between sensitivity and specificity, and automated contamination cleanup should prioritize retaining correct sequences in genomes. Simulated experiments suggest that FCS-GX has a low false positive rate that is generally robust to varying species relatedness in the reference database. To further account for potential false positives in prokaryote genomes, we currently do not flag prokaryote-in-prokaryote contamination for removal if below 1% of total genome length. Within eukaryotes, while the low rate of contamination (0.003%) in current RefSeq genomes further supports high specificity, we are continuing to explore potential false positives especially for intra-Metazoan contamination, where 90% of the identified sequences are below 1 kb (median 279 bp).

Contamination detection is not a solved problem and the researcher’s needs will determine the best approach given the various tools’ strengths and weaknesses [2]. FCS-GX is fast and has high specificity but hasn’t been developed to characterize contaminants at lower taxonomic ranks, such as prokaryote sequences below the family level that might be present in multiisolates, MAGs, or uncultured samples. Also, by favoring higher fidelity and lower compute cost, FCS-GX does not automatically incorporate secondary data types (e.g., GC content, sequencing read depth) that may provide additional support for contaminant identification. Despite the increasing number of contamination detection tools, there have been few comparative analyses across several tools [2, 27, 28] owing primarily to differences in scope and features for providing contamination information. Careful attention is needed when considering the union or intersection of multiple methods, with the former minimizing false negatives but increasing noise and the latter minimizing false positives but decreasing sensitivity.

We are phasing in FCS-GX screening of new genomes submitted to NCBI with reporting and removal of most contaminant calls starting in May 2023. As a publicly available tool, FCS-GX can be used to detect contaminants in draft or proprietary genome assemblies, minimize potential errors in analyses, and streamline later submission. While contaminant sequences can lead to errors in interpretation, they may represent novel species and themselves be of scientific value. Based on user input, we’ve included an option to bin contaminant sequences into a separate file for further review, which can be submitted as a separate metagenome or MAG assembly. The archival nature of the INSDC databases means existing contamination cannot be readily removed without the cooperation of the original data submitters. Comprehensive reports including all assemblies screened to date are available on NCBI’s genomes FTP site, and we are developing methods to provide FCS-GX contamination reports with individual genomes that users can use to filter out genomes or hardmask sequences, and exploring options for labeling sequences and reducing contaminants in BLAST databases.

## Conclusions

FCS-GX facilitates the rapid identification and removal of contaminant sequences from assembled genomes of both eukaryotes and prokaryotes, enabling assembly providers to improve data quality and avoid artifacts that impact downstream analyses. We measured FCS-GX specificity >99.93% with artificially fragmented genomes, and sensitivity >95% for many species of contaminants. Our analyses of over 1.6 million assemblies show genome contamination is present in many current genomes to varying degrees, with more than half contributed by 161 egregious cases, and we used FCS-GX to improve RefSeq to only 0.01% contaminant bases. We encourage widespread adoption of FCS-GX to improve quality of the ongoing explosion of new genomes, in particular by groups generating assemblies, building an assembly resource, or brokering the submission of assemblies to the INSDC archives. We welcome feedback from the scientific community for further improvements through our GitHub site or NCBI’s help desk.

## Methods

FCS-GX is a genome-wide contamination detection and removal tool designed for assembled sequences (**Fig. 1**). A user-supplied taxonomic identifier (tax-id) is used to distinguish contaminants from sequences corresponding to the source genome. The primary output of FCS-GX is a cleaned FASTA with its associated action report that lists sequences and sequence ranges identified as contaminants alongside executed actions to clean contamination from the genome. FCS-GX is written in C++ and Python and is deployed as Docker and Singularity containers.

### Construction of the FCS-GX database

FCS-GX uses a reference database of sequences hosted at NCBI to enable contamination detection in new genome assemblies. The FCS-GX database version r2023-01-24 contains sequences from 28,213 RefSeq and 19,541 GenBank assemblies totaling 709 Gbp of sequence data. The database contains higher counts of genomes from prokaryotes and eukaryotes with small genomes which are more likely to be genome contaminants and lower counts of eukaryotes with large genomes to avoid overinflating the size of the reference database. To select genomes for taxonomic groups, we prioritize reference genomes from highly researched taxa based on the significance criteria in NCBI Assembly, which factors in counts of NCBI BioProject submitters for the organism. We select additional representatives to achieve sufficient taxonomic diversity while maintaining a target database footprint of ∼470 GiB. We calculate Jaccard distances between the proteomes of assembly pairs and successively add assemblies that are taxonomically distinct. We exclude sequences <10 kbp in eukaryotic assemblies and <1 kbp in prokaryotic assemblies as these are more likely to contain contaminants. We progressively identified likely contaminants using FCS-GX with a leave-one-out screening approach through multiple iterations of database construction, and excluded contaminant sequences in the database using bulk analysis of FCS-GX results, a manually curated exclusion list, as well as excluding entire assemblies with >20% by length assigned as contaminant. The public FCS-GX code also includes an exclusion list which is used to further refine results, allowing newly identified contaminants in the database to be dropped with a new software version without requiring a full rebuild of the database.

FCS-GX uses a locality-sensitive-hashing approach [29] to index reference sequences for subsequent alignment steps. We extract 56-mers from sequences and drop every third base, which corresponds to codon-wobble positions if the h-mer is in a coding region and in-frame. The h-mers are collected from sequences with a 10 bp stride for prokaryotes and 20 bp stride for eukaryotes to cover all reading frames. We transform the two-bit nucleotide alphabet {A, C, G, T} to one-bit {[AG], [CT]} since transition-type mismatches are generated at a higher frequency relative to transversions. We perform a minword function min(h(k-mer), h(reverse-complement(k-mer))) to make the hash value invariant with forward or reverse sequence orientation. We store the final 38-bit h-mers index with the 2-bit-coding nucleotide sequences of the database subjects and metadata files on a RAM-backed filesystem to enable parallelization over multiple CPUs.

### FCS-GX alignment

#### Sequence pre-processing

Contaminants can be assembled as contigs within larger scaffolds. We split query sequences on runs of Ns of 10 bp or longer that are commonly used to delimit contigs within scaffolded sequences. We align each split sequence to the FCS-GX database. In order to cap the working-memory usage and improve parallelization, we split sequences into 100 kbp chunks with a 100 bp overlap and align each chunk separately in parallel. The alignments from different chunks are subsequently combined as if the sequence was aligned whole.

Repetitive regions can generate noise when assigning sequences to taxonomic divisions. We detect transposon-like sequences in eukaryotes by computing h-mer statistics and mask putative transposons as overrepresented h-mers. In order to differentiate real transposons from egregious contamination by a large number of short similar sequences (e.g. a viral infection or phiX174), the scope of h-mer statistics is limited to genomic sequences of length at least min(100kb, sequence N80 before splitting on Ns). Transposon masking is not used in prokaryote and virus genomes. We mask the low-complexity regions with a DUST-like algorithm [30] by scanning over the sequence with a 50 bp sliding window and identifying regions where the Shannon entropy of distribution of hexamers in the window falls below a threshold value of 4.5.

#### Alignment

In the alignment stage, we preprocess initial seeds to identify regions of potential homology. For every position in the query sequence, we locate the set of subjects-positions of the corresponding h-mer in the database. The tuples (query-pos q, subject-seq-id, subject-pos s) form the set of initial seeds for downstream alignment refinement. The coordinates q and s are signed indicating strand orientation on the query relative to the subject. To remove spurious matches, we apply a noise-filtering step to keep only those seeds having a close neighbor within 1 kbp on the diagonal and within 10 kbp on the antidiagonal:

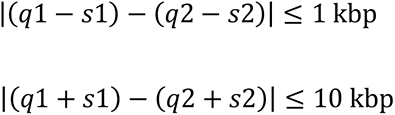

We filter the resulting set of alignments on a per-taxon basis to keep only the best alignments among those that overlap on query coordinates.

To improve the database coverage estimations on query sequences, we conduct a second alignment pass using select taxa from the FCS-GX database identified from the first alignment pass. For each sequence, we calculate the number of ungapped alignments, maximum alignment length, summed alignment length, and squared summed alignment length for all taxa from the database with at least one match. Taxa placed in the top three of any of these metrics are retained for the second alignment pass. We build a new sequence index from query sequences on-the-fly using smaller h-mers (20 bits) and align the subject-neighborhoods (limited to min(100Kbp, 2*query-length)) for the selected taxa against the query-index. We refine local alignments by first performing ungapped extensions on the seeds, followed by gapped extensions, repeatedly finding an indel that yields highest-identity continuation-seed near the alignment’s end and extending that, until the extension no longer yields a significant ungapped segment.

Alignment scoring methods that use a weighted sum of matches and mismatches (e.g., BLAST [31, 32]) do not always take into account the distribution of mismatches. Alignments with fewer mismatches that are more broadly distributed across an alignment often represent a taxonomically distant alignment relative to alignments with more mismatches that are clustered, the latter of which can arise from alignment artifacts, sequencing artifacts, or multi-nucleotide mutations. As such, we define the score of a segment as square root of sum of squares of alignment lengths with 100% nucleotide identity.

### Taxonomic assignment

FCS-GX reports alignment information including alignment coverage length and score for up to a maximum of four species per sequence. We use tax-ids to categorize species into taxonomic divisions following aggregation and modification of BLAST divisions. We limit reporting for up to a maximum of two species per division per sequence, as this can help distinguish valid contaminants from false positives arising from contamination in the database. We group taxonomic divisions into six broader kingdoms: animals (Metazoa), plants (Viridiplantae), fungi, protists (other Eukaryota), Bacteria, and Archaea. FCS-GX processes alignment coverage and score values to generate taxonomic assignments and compares assignments to the user-supplied tax-id corresponding to the source organism. Each sequence is placed into one of three broad categories: primary division (taxonomic division is consistent with input tax-id), contaminant (division is different), or inconclusive (not enough information or conflicting information from multiple divisions prevents a conclusive taxonomic assignment).

In certain cases, the sequences from a single organism may have top scoring alignments to multiple taxonomic divisions, and these hits should collectively be treated as belonging to the primary division. For all taxonomic divisions with sequences assigned to that division, we calculate the degree to which the alignments for each division overlap the alignments corresponding to the division with the highest coverage. Divisions with high overlap percentages from the same taxonomic kingdom are assigned as a set of inferred primary divisions that FCS-GX treats as belonging to the declared source organism. The degree of concordance to which the alignments must overlap is dynamic and is lower for poorly-represented species.

When using reference-based contamination detection methods, a low proportion of sequences assigned to the primary division could be the result of (a) misclassified genomes in the reference database, (b) high contamination level, (c) user error specifying an incorrect tax-id, or (d) rare genomes with poor taxonomic representation in the reference database [2]. As mentioned above, FCS-GX has a reporting strategy to include multiple species and taxonomic divisions which can reduce errors occurring from (a). In rare cases of (b) or (c), the user-asserted source organism would not be in the inferred primary division set. If a user-asserted division is well-represented in the FCS-GX database and the assembly has poor coverage from that division, then the division(s) calculated as the primary set are instead reported as contaminants along with a user warning. To reduce the incidence of false positive contaminant calls from (d), FCS-GX calculates the aggregate coverage across the genome by dividing the total sequence length aligned to the top four hits by the total genome size and applies a minimum coverage cutoff threshold for calling contaminants. Thus, FCS-GX restricts contaminant calls to the highest confidence calls in genomes with fewer close taxonomic neighbors in the database.

### FCS-GX output

The primary output of FCS-GX is a cleaned FASTA and a report that lists contaminant sequence identifiers, their associated taxonomic assignments and coverage values, and one of six recommended actions for cleaning contamination from the genome assembly (**Additional file 1: Fig. S1**). Three FCS-GX actions represent high confidence contamination calls and result in automated removal of contaminant sequences during the genome cleaning step (EXCLUDE, TRIM, FIX). Sequences with high contaminant coverage that should be removed from the assembly are assigned the EXCLUDE action, and chimeric sequences with terminal or internal contamination are expected to be rare and assigned the TRIM and FIX action, respectively. Three FCS-GX actions represent lower confidence contamination calls and warrant inspection by the user but do not result in automatic removal during the genome cleaning step (INFO, REVIEW, REVIEW_RARE). Chimeras involving sequences that are known to be integrated into host genomes (e.g., bacterial endosymbionts) are assigned the INFO action. Intra-kingdom chimera and sequences with low contaminant coverage from a division with evidence of contaminant sequences elsewhere in the genome are assigned the REVIEW action. Small amounts of prokaryote-in-prokaryote contamination (<1% of the genome assembly size) are assigned the REVIEW_RARE action.

### Computing sensitivity and specificity

We retrieved metadata for 663 prokaryote and 370 eukaryote genome assemblies for sensitivity and specificity testing using eutils v.19.0 [33]. We restricted results to NCBI non-reference assemblies with a contig N50 ≥100 kbp and ≥1 Mbp contiguous sequence and filtered out partial and anomalous assemblies. We cross-referenced the result set against the FCS-GX reference database and selected cases having the same species in the database but removed cases that were already present in the database. We randomly selected a single assembly per genus to test. For each assembly, we retained sequences ≥1 Mbp using seqkit v.0.11.0 [34] as these sequences are more likely to be contaminant-free. Following manual inspection, we removed eukaryotic assemblies with sequences >1 Mbp assigned as prokaryote from the analysis. After splitting scaffolds on runs of 10 Ns with FCS-GX, we applied a second size-selection filter to remove contigs <100 kbp. We split the remaining contigs three separate times into subsequences of defined sizes (1 kbp, 10 kbp, 100 kbp) using seqkit. We ran two separate FCS-GX pipelines on split sequences: we supplied a false tax-id with the input FASTA to test sensitivity and we supplied the true tax-id with the same FASTA to test specificity. Prokaryote sensitivity tests used the tax-id for human (NCBI:txid9606) and eukaryote sensitivity tests used the tax-id for *E. coli* (NCBI:txid562). We turned off FCS-GX repeat masking for sensitivity tests in both prokaryotes and eukaryotes. We ran the same sequence sets while excluding alignments to the species tax-ids of the source genome to simulate contamination detection in a novel organism.

We measured sensitivity as the percentage of sequences assigned as contaminant with FCS-GX corrective actions (EXCLUDE, TRIM, FIX) corresponding to the taxonomic division of the source genome. We considered sequences assigned as prokaryote virus to be true positives when measuring sensitivity in prokaryote genomes. We measured specificity as the percentage of sequences assigned as non-contaminant by subtracting contaminant sequences assigned a FCS-GX corrective or review action (EXCLUDE, TRIM, FIX, REVIEW) from the total number of sequences. To measure sequence-level specificity (**Additional file 2: Table S4**), we estimated an upper bound using the equation 100*(1 - sum(lengths of all same-kingdom contaminant calls in all genomes)/(sum(lengths of all genomes) - sum(lengths of all cross-kingdom contaminants)). We estimated a lower bound using the equation 100*(1 - sum(lengths of all contaminant calls in all genomes)/(sum(lengths of all genomes)). For visualization, we separated genomes into five “kingdom” groups derived from the NCBI taxonomy [21]: prokaryotes (Bacteria and Archaea), Metazoa (animals), Viridiplantae (green plants), Fungi, and other Eukaryotes.

### GenBank and RefSeq genome datasets

Assemblies were identified for screening from the assembly_summary.txt reports available on the NCBI genomes FTP site as of 4/15/2023, selecting all eukaryote and prokaryote assemblies with available FTP files. Eukaryote assemblies were run on a Dell PowerEdge c6525 server with a 32 core Intel Xeon processor with hyperthreading enabled for 64 vCPUs and 1 TiB RAM. Assemblies were run using 16 cores each and up to 4 parallel jobs, with 74% observed CPU utilization. Prokaryote assemblies were run on available shared servers in parallel using 1-4 cores each and up to 64 cores total using xargs. Results were stored in a custom SQL database. Relative CPU usage for FCS-GX compared to megaBLAST was estimated from aggregate data on submissions over six month windows and normalized to total genome file size, which approximates submitted genome size. The set of assemblies available at different points in time were determined based on assembly release date and not-live date, which is the point at which a given assembly version was last replaced or suppressed.

GX results were compared to Conterminator based on the published data provided at https://figshare.com/projects/Conterminator/77346. Sequence accessions were mapped to assemblies to identify the subset of assemblies with one or more reported contaminants found by Conterminator. Accessions reported by FCS-GX, Conterminator, or both were then quantified.

### Cleaning RefSeq genomes

GX results were reviewed for RefSeq genomes with higher levels of contamination using development versions of FCS-GX and the FCS-GX database. In some cases, additional contaminant sequences were identified from FCS-GX intermediate results, or through BLAST or other datasets. False positives were identified and excluded from cleanup. The original genome submitters were consulted, and with their permission contaminant sequences were suppressed and new versions of both the GenBank GCA_ and RefSeq GCF_ assembly were released with a revised assembly name and a public comment about the change. Chimeric sequences are harder to address in released sequences and were only revised in significant cases such as sperm whale chromosome 1 CM014785.2 / NC_041214.2. For genomes originally submitted through ENA or where the submitter could not be reached, only RefSeq sequences were suppressed, but no sequences were changed. For prokaryotes, the primary action taken was to mark assemblies as ‘contaminated’ and exclude them entirely from the RefSeq collection.

### Lateral gene transfer analysis

We downloaded the metadata file HGT_data.txt.zip from the mobile genetic elements resource https://promge.embl.de/ reported in Khedkar et al. [24]. We used the sequence identifiers listed in the genomic coordinates column to retrieve a set of assemblies with putative LGT. We cross-referenced this set against the set of GenBank assemblies with FCS-GX results produced in this study (n=3,986). We determined the frequency of chimeric FCS-GX calls (TRIM, FIX, REVIEW or REVIEW_RARE if start_pos + end_pos != length) in both the LGT set and all current GenBank assemblies. We compared the coordinates of FCS-GX chimeric calls to the coordinates reported in Khedkar et al. and used BLAST to inspect cases where coordinates overlap.

## Declarations

### Ethics approval and consent to participate

Not applicable

### Consent for publication

Not applicable

### Availability of data and materials

The datasets generated and/or analyzed during the current study are available at the NCBI FTP site https://ftp.ncbi.nih.gov/genomes/TOOLS/FCS/reports/20230416/ [35]. FCS-GX code is written in Python and C++ and available at https://github.com/ncbi/fcs-gx [36]. The static version of FCS-GX used in this work (v0.4.0) is available at https://github.com/ncbi/fcs-gx/releases/tag/v0.4.0. FCS-GX is part of the NCBI Foreign Contamination tool suite available at https://github.com/ncbi/fcs [37].

### Authors’ contributions

A.A., T.D.M. conceived the project. A.A. developed core FCS-GX algorithms. D.S., N.B., L.W., V.S., V.J., and P.M.S. wrote code for FCS-GX algorithms and infrastructure. V.S. and E.M. implemented FCS-GX in NCBI submission screening. E.S.T., P.K.S., B.S.W., S.L.B., K.C., and T.D.M. analyzed the data. E.S.T. and T.D.M. prepared data visualizations. E.S.T., A.A., and T.D.M. wrote the original manuscript. All authors read and approved the final manuscript. W.H., E.W.D., V.A.S., K.D.P., and T.D.M supervised the project.

## Acknowledgements

The legacy contamination screening process at NCBI was conceived by Paul Kitts. We acknowledge Hena Bajwa, Eric Cox, Stacy Cuifo, Mike DiCuccio, Jinna Hoffman, Brad Holmes, Anne Ketter, Avi Kimchi, Peter Meric, Barbara Robbertse, Guangfeng Song, Françoise Thibaud-Nissen, Jovany Tinne, and Igor Tolstoy for internal discussions and review of FCS-GX data. We further thank Therese Catanach (Drexel University), Peter Ebert (Heinrich Heine University), Natalia Ivanova (JGI), Erich Jarvis (Rockefeller University), Brenda Oppert (USDA-ARS), Michael Paulini and James Torrance (Wellcome Sanger Institute), David Roos (VEuPathDB), Vasily Sitnik (EMBL-EBI), and many GenBank genome submitters for consults on contamination results and feedback on FCS-GX development.

## Supplementary Information

### Additional file 1: supplementary figures

**Fig. S1**

FCS-GX commands and sample output.

**Fig. S2**

Summary of FCS-GX results for false negatives in sensitivity tests. For 1 kbp sequence sets, aggregate counts of false negatives are shown for FCS-GX runs while including the same species tax-ids as the source genome during the alignment stage (+species) and while excluding same species tax-ids (- species). Categories are classified as follows: Review – sequences with the FCS-GX action REVIEW that are assigned the proper contaminant taxonomy but with subthreshold alignment coverage, Virus – sequences assigned prokaryote virus in prokaryote genomes and eukaryote virus in eukaryote genomes, Contaminant (inter-kingdom) – sequences assigned as contaminant by FCS-GX but the taxonomic classification is wrong and is in a different kingdom grouping. Contaminant (intra-kingdom) – sequences assigned as contaminant by FCS-GX but the taxonomic classification is wrong and is the same kingdom grouping. Non-contaminant – sequences assigned as non-contaminant. See **Additional file 2: Table S2** for counts/percentages of all false negative categories.

**Fig. S3**

Plots of aggregate FCS-GX alignment coverage against sensitivity. Aggregate coverage is calculated as the total percentage of the genome with overlaps from sequences in the FCS-GX reference database. Results are shown for 1 kbp sequence sets for FCS-GX runs while including the same species tax-ids as the source genome during the alignment stage (blue circles) and while excluding same species tax-ids (red triangles).

**Fig. S4**

Complete distributions of specificity measurements. Distributions are shown for artificially fragmented genomes in five “kingdom” groups. Specificity is shown for genomes fragmented at three different window sizes (1 kbp, 10 kbp, 100 kbp). For each window size, specificity is shown for FCS-GX runs while including the same species tax-ids as the source genome during the alignment stage (+species) and while excluding same species tax-ids (-species). Red arrows point to ten outliers that are not visualized in Fig. 2B.

**Fig. S5**

Length distribution of contaminants detected by FCS-GX.

### Additional file 2: supplementary tables

**Table S1**

FCS-GX sensitivity and specificity scores on artificially fragmented genomes.

**Table S2**

Counts and percentages of FCS-GX false negative types for sensitivity tests on 1 kbp sequences.

**Table S3**

FCS-GX contamination calls in 100 kbp fragmented genome datasets produced during specificity tests.

**Table S4**

Upper and lower bound estimates of FCS-GX sequence-level specificity.

**Table S5**

Aggregate contamination identified in current GenBank genomes by FCS-GX, grouped by taxonomic kingdom.

**Table S6**

Aggregate contamination identified by FCS-GX contained in GenBank and RefSeq databases from 2017-2023.

**Table S7**

Aggregate contamination identified in current GenBank genomes by FCS-GX, grouped by FCS-GX taxonomic division.

**Table S8**

*Cladocopium goreaui*-like contaminants identified in GenBank genomes.

**Table S9**

GenBank genomes with extreme contamination by aggregate contamination length, representing 50% of total contamination identified by FCS-GX.

**Table S10**

GenBank genomes with extreme contamination by proportion of total sequence length (90-100% contaminated).

**Table S11**

Summary of genomes with species assignment issues identified by FCS-GX.

**Table S12**

Comparison of contaminants reported in FCS-GX vs Conterminator.

Table S13

Summary of cleaned RefSeq and RefSeq candidate genomes.

**Table S14**

Aggregate contamination identified in current RefSeq genomes by FCS-GX, grouped by taxonomic kingdom.

**Table S15**

FCS-GX chimeric contamination calls in suspected LGT regions.

