## Supplementary material for "Rapid and sensitive detection of genome contamination at scale with FCS-GX": Fig. S1

FCS-GX command for Toxoplasma gondii ME49 (GCA_000006565.2)

```
python3 ./fcs.py screen genome --fasta ./gx_in/GCA_000006565.2_TGA4_genomic.fna.gz
--out-dir ./gx_out/ --gx-db "$GXDB_LOC/gxdb" --tax-id 508771
```

Summary contamination report for Toxoplasma gondii ME49 (GCA_000006565.2)

```
fcs_gx_report.txt contamination summary:
----------------------------------------
                                seqs      bases
                               ----- ----------
TOTAL                            321     226690
-----                          ----- ----------
anml:primates                    282     169672
anml:nematodes                    34      55025
virs:eukaryotic viruses            3        819
anml:rodents                       1        639
fung:basidiomycetes                1        535

-----------------------------------------------------------------------------

fcs_gx_report.txt action summary:
---------------------------------
                                seqs      bases
                               ----- ----------
TOTAL                            321     226690
-----                          ----- ----------
EXCLUDE                          321     226690
```

Sample FCS-GX contamination action report for Toxoplasma gondii ME49

```
#seq_id start_pos      end_pos seq_len action  div      agg_cont_cov   top_tax_name
KE138932.1     1       3819    3819    EXCLUDE anml:nematodes 100     Brugia malayi
KE138975.1     1       2274    2274    EXCLUDE anml:nematodes 100     Brugia malayi
KE138976.1     1       1893    1893    EXCLUDE anml:nematodes 100     Brugia malayi
KE139040.1     1       2152    2152    EXCLUDE anml:nematodes 100     Brugia malayi
KE139041.1     1       2186    2186    EXCLUDE anml:nematodes 100     Brugia malayi
KE139042.1     1       2219    2219    EXCLUDE anml:nematodes 100     Brugia malayi
KE139043.1     1       1975    1975    EXCLUDE anml:nematodes 100     Brugia malayi
KE139071.1     1       1494    1494    EXCLUDE anml:primates  100     Homo sapiens
KE139077.1     1       1328    1328    EXCLUDE anml:nematodes 81      Brugia malayi
KE139078.1     1       1466    1466    EXCLUDE anml:nematodes 100     Brugia malayi
KE139079.1     1       1557    1557    EXCLUDE anml:nematodes 100     Brugia malayi
KE139081.1     1       1439    1439    EXCLUDE anml:nematodes 97      Brugia malayi
KE139286.1     1       1949    1949    EXCLUDE anml:primates  100     Homo sapiens
KE139334.1     1       1575    1575    EXCLUDE anml:nematodes 100     Brugia malayi
KE139335.1     1       1570    1570    EXCLUDE anml:nematodes 100     Brugia malayi
KE139336.1     1       1740    1740    EXCLUDE anml:nematodes 97      Brugia malayi
KE139337.1     1       1512    1512    EXCLUDE anml:nematodes 100     Brugia malayi
KE139338.1     1       1771    1771    EXCLUDE anml:nematodes 100     Brugia malayi
```

FCS-GX command for removing contaminant sequences (genome cleaning)

```
zcat GCA_000006565.2_TGA4_genomic.fna.gz | python3 ./fcs.py clean genome
--action-report ./gx_out/GCA_000006565.2_TGA4_genomic.fna.508771.6973.fcs_gx_report.txt
--output clean.fasta --contam-fasta-out contam.fasta
```
