## Supplementary figures and images for "Rapid and sensitive detection of genome contamination at scale with FCS-GX"

### Fig. S2

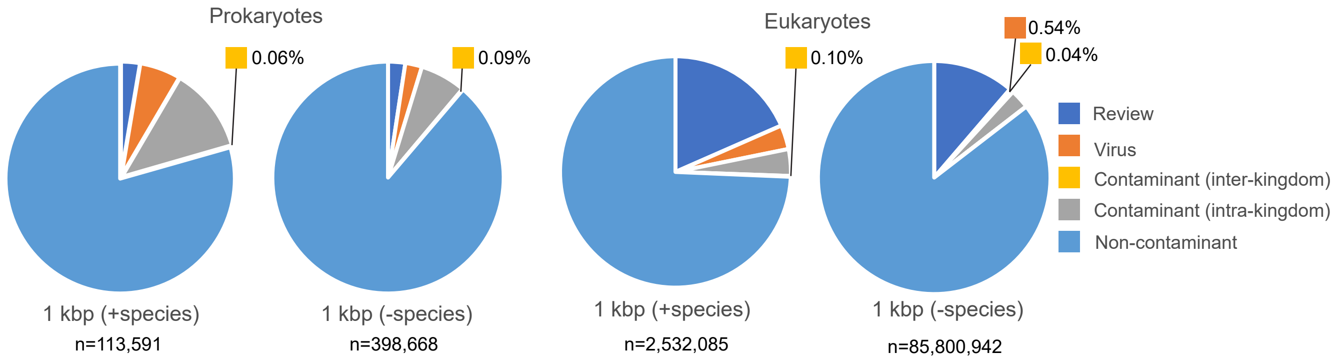

### Fig. S3

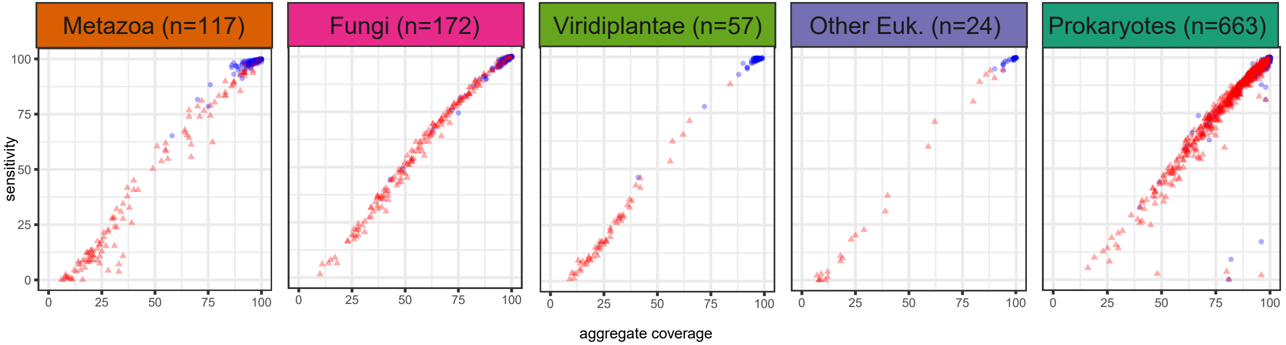

### Fig. S4

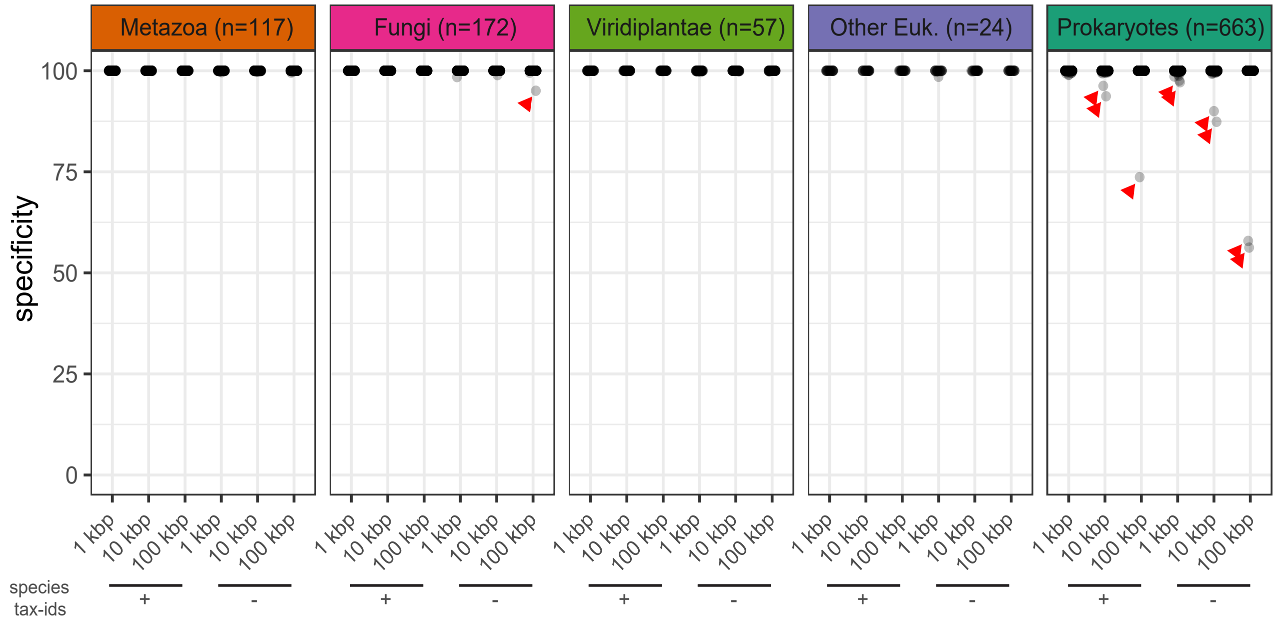

### Fig. S5

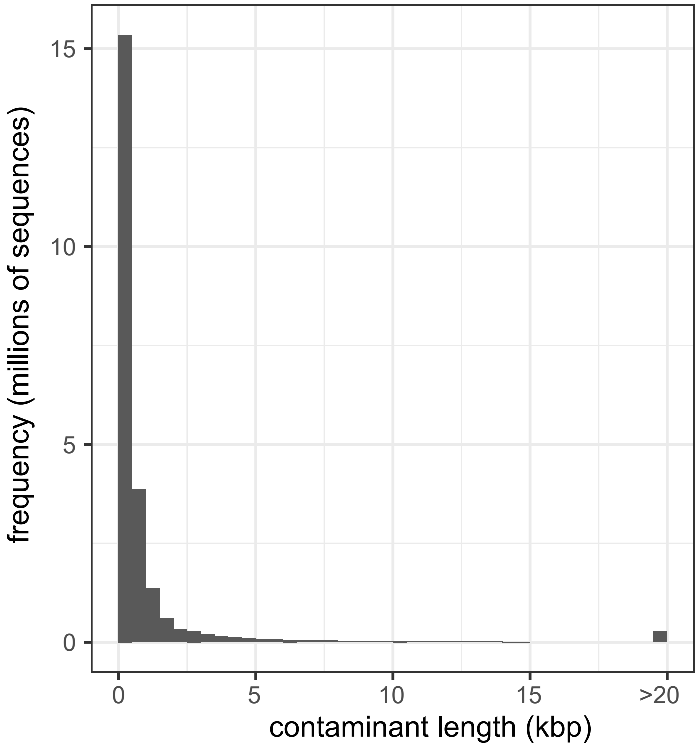
